# 4D reconstruction of developmental trajectories using spherical harmonics

**DOI:** 10.1101/2021.12.16.472948

**Authors:** Giovanni Dalmasso, Marco Musy, Martina Niksic, Alexandre Robert-Moreno, Claudio Badía-Careaga, Juan J. Sanz-Ezquerro, James Sharpe

**Affiliations:** European Molecular Biology Laboratory (EMBL-Barcelona), Barcelona, Spain; Centre for Genomic Regulation (CRG), Barcelona, Spain; Centro Nacional de Investigaciones Cardiovasculares (CNIC), Madrid, Spain; Centro Nacional de Biotecnologia (CSIC Madrid), Madrid, Spain; Institució Catalana de Recerca I Estudis Avançats (ICREA), Barcelona, Spain

**Keywords:** Spherical Harmonics, Mouse Development, Limb Growth, Heart Growth, Morphogenesis, OPT, Voxel Data

## Abstract

Although the full embryonic development of species such as Drosophila and zebrafish can be 3D imaged in real time, this is not true for mammalian organs, as normal organogenesis cannot be recapitulated *in vitro*. Currently available 3D data is therefore *ex vivo* images which provide only a snap shot of development at discrete moments in time. Here we propose a computer-based approach to recreate the continuous evolution in time and space of developmental stages from 3D volumetric images. Our method uses the mathematical approach of spherical harmonics to re-map discrete shape data into a space in which facilitates a smooth interpolation over time. We tested our approach on mouse limb buds (from E10 to E12.5) and embryonic hearts (from 10 to 29 somites). A key advantage of the method is that the resulting 4D trajectory takes advantage of all the available data (i.e. it is not dominated by the choice of a few “ideal” images), while also being able to interpolate well through time intervals for which there is little or no data. This method not only provides a quantitative basis for validating predictive models, but it also increases our understanding of morphogenetic processes. We believe this is the first data-driven quantitative 4D description of limb morphogenesis.

## 1 Introduction

Progress in imaging technology has been central to understanding morphogenesis, which is an intrinsically 3D and dynamical process. In the case of externally developing organisms, such as *Drosophila* and zebrafish, it is now possible to image *in vivo* whole-embryo development at a cellular level, up to and including the later organ-forming stages (Tomer et al. 2012, Royer et al. 2016). However, for more complex animal models (e.g. mouse embryogenesis), it is still not possible to observe organogenesis in real time, due to the limitations of *in vitro* culture techniques. The dynamics of early post-implantation mouse embryogenesis have been successfully imaged in great detail (McDole et al. 2018), however embryo culture beyond E10.5 is not robust enough to recapitulate the full development of complex organs - largely due to the lack of blood flow. *In vivo/in utero* imaging techniques (such as MRI) do not yet provide sufficient spatial resolution to capture the morphogenesis of organs, and our understanding of complex mammalian organogenesis is therefore largely derived from capturing series of *ex vivo* and static 3D images, often using mesoscopic imaging techniques such as optical projection tomography (Sharpe et al. 2002) (Figure 1A), light sheet imaging (Huisken et al. 2004), or *ex vivo* MRI (Wong et al. 2012, 2014).

**Figure 1:**
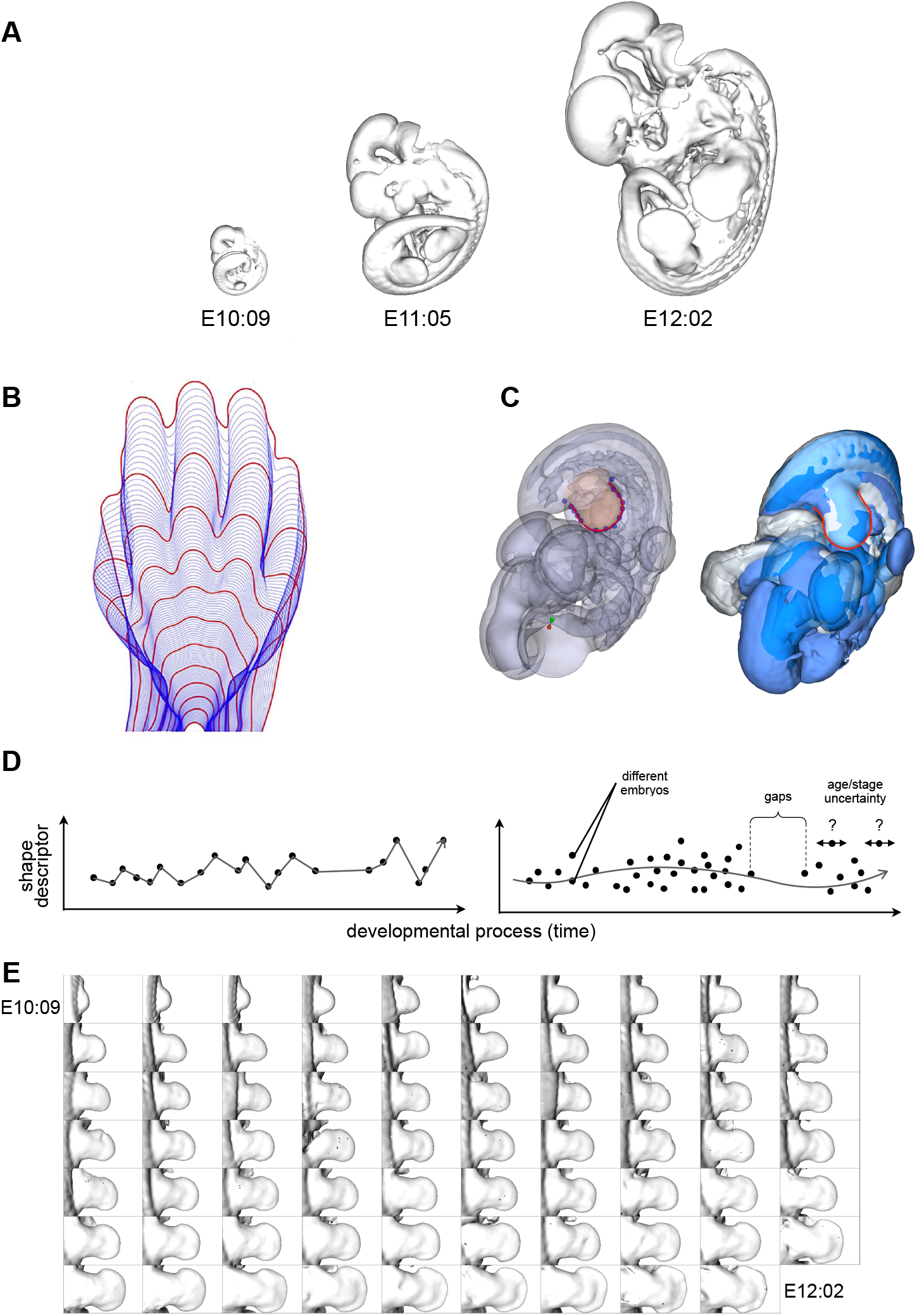
**A** - Three mouse embryos at different developmental stages (E10:09, E11:05, E12:02). **B** - Two dimensional interpolation of one mouse limb reference shape to the next in the time sequence. The red lines indicate the shapes that correspond to the experimental averages for each age group (*from* Musy et al. (2018)). **C** - Alignment of three embryos at the same developmental stage according to their right hind limbs shown with transparency (left) and as surface in solid color (right). The red line represents the output of the staging system (Musy et al. 2018). **D** - Graphical illustration of a naive linear based interpolation (left plot) versus a smooth trajectory of the data set (right plot). **E** - Data set of mouse right hind limbs used in this study ordered according to their developmental stage from E10:09 to E12:02.

An important remaining challenge is to integrate these series of static 3D data-sets into a smooth, continuous and 4D trajectory that realistically models the morphogenesis of the organ in question. There have been diverse attempts to recapitulate the development of a full mouse embryo. Among others, Wong et al. (2015), used 3D OPT images taken at six developmental time points (between E11.5 and E14.0), to create a 4D model of mouse embryo growth. Specifically, the outcome was the result of spline fitting and interpolating over time the displacement vectors of homologous points of the 3D voxel images. While this model provided a global overview of mouse embryo development, it was not able to accurately recreate smooth shape trajectories for all the organs. In particular, registration of the voxel data was not able to align parts of the embryo whose positions are intrinsically variable over time, such as the limbs and the tail. Therefore, this method was not able to precisely reproduce the development of these particular mouse’s structures (Wong et al. 2015). Indeed, the large degree of natural variation in samples represents indeed a key challenge in the effort of recapitulating the growth of a mouse embryo or even a sub-part of it such as the limb.

Two embryos at the exact same age do not look identical, due to the intrinsic variation both in shape and in development. We therefore ideally need a technique which, on the one hand can take advantage of as much shape data as possible, while at the same time not giving excessive weight to any specific individual. In practice this means a procedure in which each sample (i.e. each real limb) is able to influence the mean trajectory at ages both younger and older than itself. Or in other words, the final reconstruction needs to capture the right balance between the overall shape trends across the whole data-set, and the actual details of each limb. Another challenge is that the shape complexity of an organ increases during development. We need a method that performs some degree of dimensionality reduction, but without losing the more complex features of the older stages.

Spherical harmonics have been known since 1782 (Laplace - Mécanique Céleste) and form a natural basis for describing how a scalar quantity varies on the surface of a sphere. They produce an orthogonal basis for mathematically describing a 3D shape and provide a compact parametric representation of it. Spherical harmonics have been widely used in different fields and, in recent years, also in biology. One of their main use has been to characterize, among others, the shapes of cells and organs (Styner et al. 2006, Morishita et al. 2017, Medyukhina et al. 2020), deformations and movements of tissues in an easier and more robust way (Khairy et al. 2018). Moreover, spherical harmonics expansion has been also applied to mesh refinements (Lai et al. 2009). The expansion of a function in spherical harmonics is characterized by a finite set of coefficients which multiply the set of functions constituting the orthogonal basis. These coefficients encode the information of the original function, therefore, the higher the order (i.e. the number of coefficients used) the better the representation. Among other properties, one important feature that makes spherical harmonics particularly convenient is that the information encoded in the coefficients decreases with their order. Therefore, limiting the number of coefficients of the series will not cause a loss of the main characteristic of the shape represented. Previous studies using spherical harmonics have considered collections of objects (both surfaces or volumes) often analysed for the purpose of quantifying the differences or similarities between the sets. We therefore propose and demonstrate an approach to describe the evolution in time and space of mouse limbs both from volumetric and surface data. We are only aware of one previous use of spherical harmonics to characterise a changing morphology over time - in the chick embryo (Morishita et al. 2017), and believe this is the first application to a mammalian organ.

A 2D trajectory of limb bud shape change has previous been created (not using spherical harmonics), and has been developed into a method to estimate the embryonic stage of a limb bud to a high temporal resolution (Musy et al. (2018), Figure 1B). This work, although only in 2D, allows us to have a reference to align in space and time mouse embryos according to a chosen limb (Figure 1C, E). Here we develop and demonstrate two different versions of this approach for the case of limb development. We believe this approach has widespread applicability in developmental biology, to provide a quantitatively reliable baseline description of organ development - something surprisingly absent in the field so far - and so we also demonstrate its use on 3D images of another mouse organ, the developing heart.

## 2 Results

One key step in spherical harmonics expansion is to first map a surface onto a sphere. In the case of relatively simple surfaces, one possible solution is using as mapping the geometrical distance between the surface considered and a sphere which contains it. Specifically, all these distances represent a set of scalars onto the sphere which serve as a mapping that can subsequently be expanded in spherical harmonic. We applied this procedure to 69 surfaces of mouse limb buds imaged using optical projection tomography (OPT) of mouse embryos previously ordered in time using the embryonic Mouse Ontogenetic Staging System (eMOSS). Expanding a function into a spherical harmonic series and reconstructing it from the spherical harmonic coefficients are the two essential procedures applied when operating with data on the sphere. However, in our work, instead of using the exact coefficients’ values of the spherical harmonic series for reconstructing the original function, we interpolate them in time. In this way, the interpolation not only provides a spatial average estimate in the points where there are more than one surface but it also produces a temporal estimate for the existent gaps due to missing data. The results is a reconstruction describing a continuous and smooth changing shape over space and time.

### 2.1 Application on simple surfaces

An arbitrary surface, which can be mapped onto a sphere, can be expanded into a finite series of spherical harmonic terms and reconstructed back. The precision by which the reconstructed surface will resemble the original one depends on the number of coefficients of the expansion, the higher the number of coefficients the more precise the reconstruction.

As a data set, we used a collection of mouse embryos polygonal surfaces extracted from OPT scans (Sharpe et al. 2002). All the embryos were staged, using the staging system already described, in order to be correctly positioned in time. We then focused on the right hind limbs obtaining a set of 69 limbs from E10:09 to E12:22 (Figure 1D).

The embryonic Mouse Ontogenetic Staging System (eMOSS) (Musy et al. 2018) provides us with a reference to align in space the limb buds. Afterwards, the alignment of the sequence of limbs was refined using the iterative closest point (ICP) algorithm (Besl & McKay 1992).

Since the limb is a rather simple surface, in order to map it onto a sphere, we used the geometrical distance between its surface and a sphere that circumscribes it (Figure 2A top-left). Along a given radius *r*_*j*_, we measured the corresponding distance *d*_*j*_ between a given surface **v** = (*x, y, z*) and the sphere. We used 500 radii along which computing these distances creating a set of 500 scalars **d** for each limb. On top on each sphere we obtain therefore a collection of sample points (equal to the number of radii) to which is associated a set of scalars **d** representing the distances between these points and the surface, as shown by the color heat-map in Figure 2A (top-right). Moreover, the number of these sample points provides the resolution of the spherical mapping of the considered surfaces. Given the total number of limb surfaces *N* we have therefore a set of scalars **d**_0_*, . . .,* **d**_*N*_ which encodes the spherical mapping of each surface. Every spherical mapping can then be expanded in spherical harmonics.

**Figure 2:**
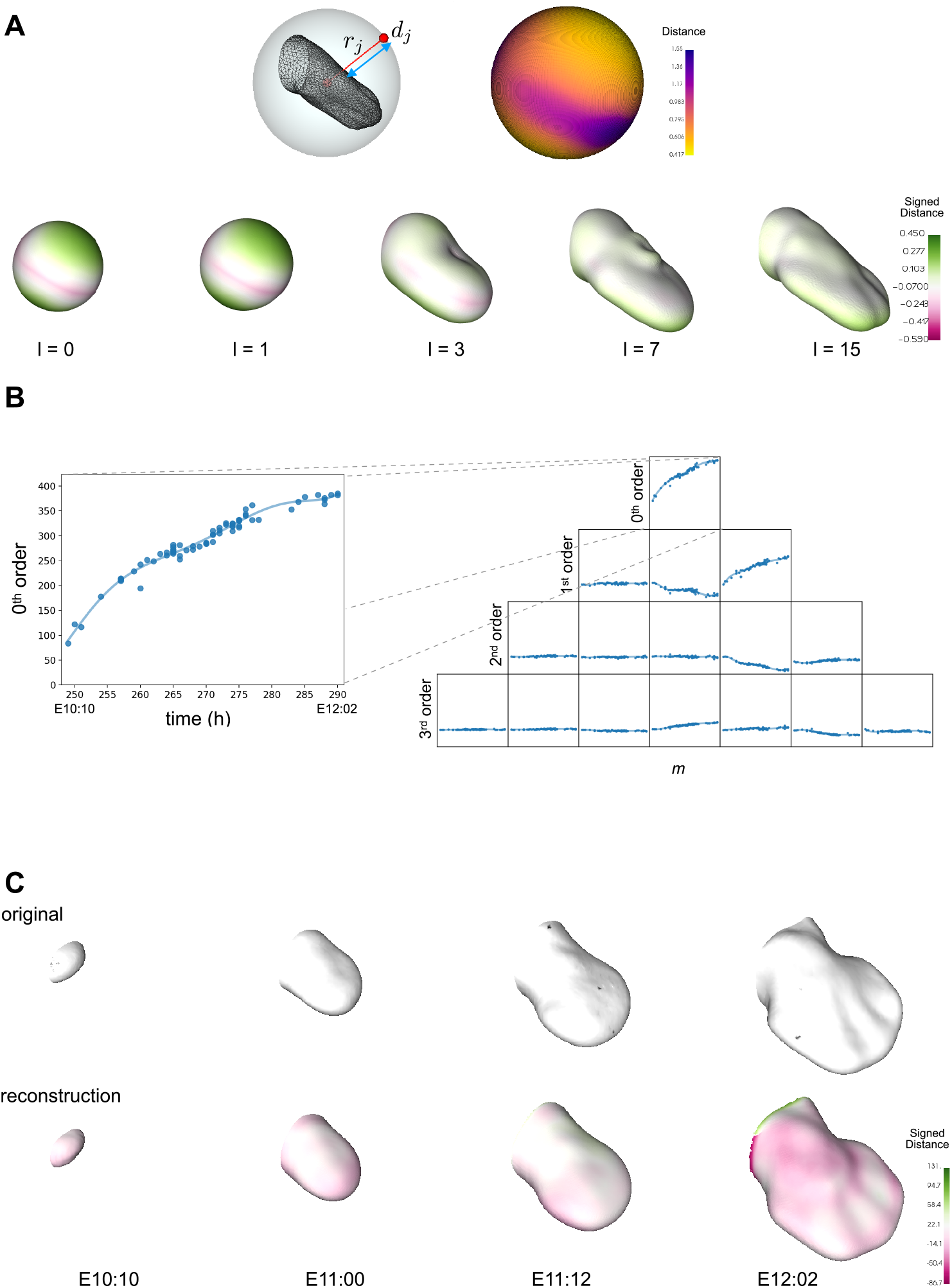
**A** - Mapping of a limb onto a sphere using geometrical distances (**d**) along the radii (**r**) between its surface and the sphere that circumscribes it (top-left). Spherical mapping of a limb, the heat-map shows the distance between the points on top of the sphere and the surface of the limb (top-right). Representation of the limb using spherical harmonic expansion of different orders, *l*_max_ = 0, 1, 3, 7, 15 (bottom), color map represents the distance between the reconstruction and the original surface. **B** - Splining in time over the spherical harmonics coefficients 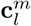. Here the first fourth orders are shown. **C** - Comparison of original data (top) at four different developmental stages (E10:10, E11:00, E11:12, E12:02) with the corresponding limb growth reconstruction (bottom), color map represents the signed distance between the reconstruction and the original surface.

A surface **v**, mapped as *x*(*ϑ, φ*), *y*(*ϑ, φ*), *z*(*ϑ, φ*), can be represented in the form:

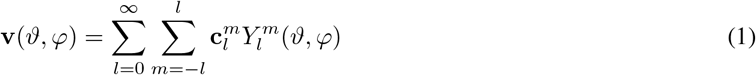

where:

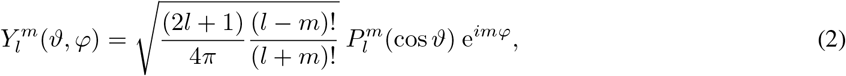

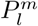 are the associated Legendre polynomials and 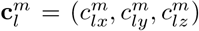, with *l* = [0, ∞) and *m* = [−*l,* +*l*], are the coefficients of the expansion. In particular, 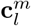 assume the form:

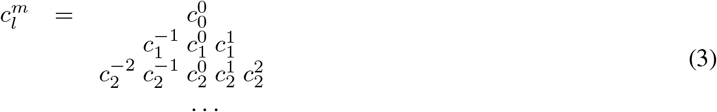

In the practical use, the coefficients 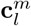 of the expansion are truncated to a specific order *l* = *l*_max_ which defines the level of detail. Spherical harmonic expansion provides a unique description of a surface based on scalar coefficients.

Using only the coefficients of the first orders (i.e. *l*_max_ = 0, 1) provides as a result practically a sphere, with *l*_max_ = 3 the details of the reconstruction increase and adding more orders (*l*_max_ ≥ 7) it is possible to obtain a limb-like surface (Figure 2A bottom). Moreover, the higher the order of the coefficient the less information it provides to the representation (i.e. in this reconstruction we are not interested in the finer morphological details). Therefore, every surface in our data set could be represented by a finite set of coefficients of its corresponding spherical harmonic expansion. This means that we have the coefficients for each surfaces at its corresponding point in time. If we put together and represent all the coefficients of e.g. order zero we will have a set of scalars evolving in time (Figure 2B left), and the same happens for the subsequent orders (Figure 2B right). Instead of reconstructing each surface using the exact coefficients we interpolated through them and used the interpolated values for the reconstruction (Figure 2C). In this way, the interpolated curve will give an average of the coefficients in the time points in which we have more than one surface and also will fill the time gaps in the points in which there are no data. Using this approach for the coefficients of all orders considered (i.e. *l*_max_ = 25) it is possible to obtain a reconstruction in space and time of the data set. In this reconstruction each limb contributes the same to the average growing shape.

As a further improvement, we used this reconstruction to refine the alignment of the original data-set obtaining a better and more precise sequence of limbs (Figure 2C top). Reapplying the method explained above in a way analogous to a bootstrapping process. With the new alignment of the data we obtain an improved reconstruction of a growing limb bud trajectory (Figure 2C bottom). To our knowledge this is the first data-driven 4D description of limb bud development across time and space (the result can only be well-appreciated from watching Movie 1 - https://vedo.embl.es/fearless/#/limb).

#### 2.1.1 Adding the limb flank

In 2.1 we were able to recreate the growth in space and time of a mouse limb. In reality however, the boundary between the limb and the rest of the embryo is rather arbitrary, and we therefore wanted to extend the analysis to include the flank of the embryo. Specifically, we extended the limb bud shape up to the mid-line of the mouse, as defined by the spinal column (Figure 3A). The new shape is now much more complex than before, displaying concavities (between the limb and the flank), and the spherical mapping described above is no longer applicable. Mapping methods for more complex surfaces and volumes have been extensively investigated (Brechbuhler et al. 1995, Shen & Makedon 2006, Lai et al. 2009, Shen et al. 2009, Yu et al. 2010), but nevertheless these proposed methods are not suitable when multiple objects need to be consistently mapped onto the same reference since e.g. the portion of the sphere of the first one may not represent the same part of the second object.

**Figure 3:**
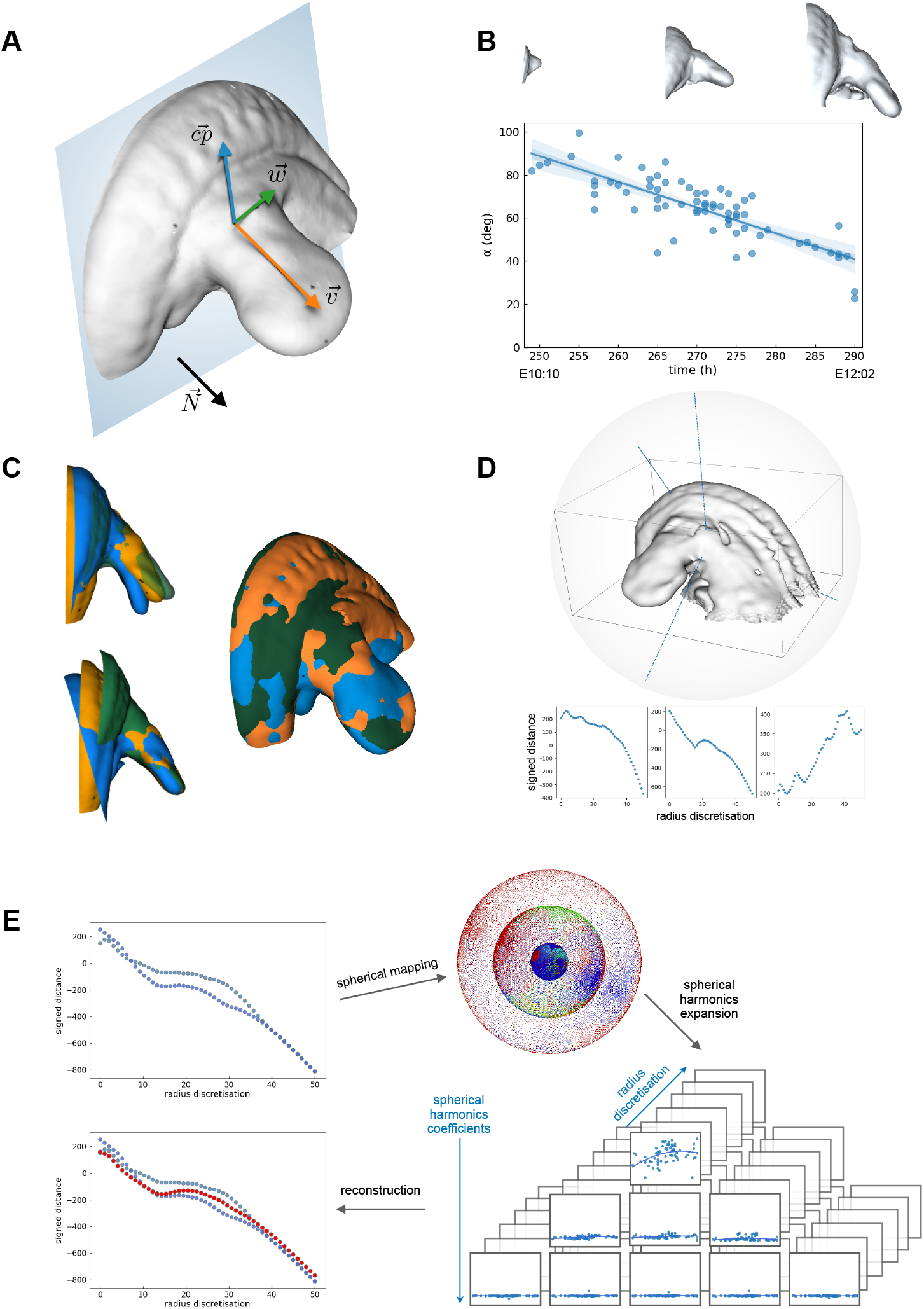
**A** - Triad 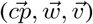 describing the orientations of the limb with respect to the normal 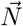 to a perpendicular plane. **B** - Variation in time of 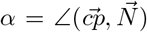 and its corresponding linear fitting with three representative limbs - E10:10, E11:25, E12:02 (top). **C** - Alignment of three limbs whit flank of the same developmental stage according to the flanks (top left) and the limbs (bottom left), alignment according to the limbs and warping of the flanks (right). **D** - Example of signed distances computed on three different radii of a limb. **E** - Example of signed distance computed on the same radius of two limbs at the same developmental stage, their spherical mapping, first three order coefficients of the spherical harmonics expansion, and comparison of the signed distance reconstruction (red dots) with the original ones (blue dots).

To map our complex embryonic surface onto a sphere, we chose to make use of volumetric data. For any arbitrary surface it is possible to embed it into a volume and to compute the signed distances over this volume from the input mesh (Bærentzen & Aanæs 2005). The output is a volumetric dataset whose voxels contain the signed distance from the mesh. Inscribing this volume into a set of concentric spheres allow us to generate a set of scalar fields (i.e. signed distances) on the sphere surface (Kazhdan et al. 2003, Skibbe et al. 2009). This allows us to expand the scalars of each sphere into spherical harmonics. In this way we are able to have a spherical harmonic representation for any kind of surfaces. It is now possible to apply this procedure on the same data set described before but we can now consider not only the limb but also the flank (i.e. part of the mouse back) since we do not have any limitation on the kind of surface to be analysed. Consequently, we can adopt the same concept used before of interpolating the spherical harmonics coefficients in time (over the developmental stages).

For a variety of reasons (external forces, natural variation, etc.) the limb buds of two embryos of the exact same age may not protrude from the flank at the same angle. But a reasonable average shape cannot be calculated from two images that do not align. To create a reliable average trajectory of normal development we need to make a judgement of which angle is the most normal, and we must then either select images that fit this average, or alter the remaining data sets so that all limb buds and all flank regions can be well-aligned. The first step was to measure the correct average angle. By defining an orthogonal triad of vectors 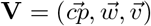 it is possible to describe the movements and rotations of the limb in space. Moreover, considering a plane perpendicular to the limb, and therefore parallel to the flank, we can compute the angles between the triad **V** and a normal 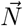 to this plane (Figure 3A). Specifically, 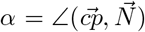 describes the vertical bending of the limb with respect to the flank and thus we can plot the angles for many embryos over time, which reveals a fairly linear decrease in this angle from about 90 degrees at mE10:10 to about 40 degrees at mE12:02 (Figure 3B). The second step was therefore to use a linear fit of *α* as a reference, and to apply a bending transform onto the shape data using *thin plate splines* (Bookstein 1989), which is a non-linear transformation based on a physical analogy involving the bending of a rigid material. The limbs were not affected by this transformation, only the non aligned flanks were bent. This operation ensures that all data-sets reflect the average bending trajectory over time (Figure 3C).

To implement the idea mentioned above, we embedded each surface into a volume and computed the signed distances over this volume from the input mesh (Bærentzen & Aanæs 2005). The output is a volumetric dataset whose voxels contain the signed distance from the mesh. It is therefore possible to generate a scalar field by the signed distance from a mesh. Inscribing this volume into a sphere it is possible to know the intensity of the signed distance (SD) along each chosen radius (*r*_0_, *r*_1_, *. . .*, *r*_*n*_) of this sphere making this mapping method theoretically applicable to any arbitrary surface (Figure 3D). Since each radius is discretized in *m* = 50 points, we have a set of *m* concentric spheres on which the signed distances are grouped according to their position along each radius. This represent a suitable condition that allows us to expand the scalars of each sphere into spherical harmonics.

With this mapping method, we obtained the 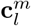 coefficients for each concentric spheres of every limb. Similar as in 2.1, instead of reconstructing each volume using the exact coefficients 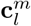 we first interpolated them over time. Subsequently we used for the reconstruction the values of the interpolation, obtaining in this way one new limb for every time point (Figure 3E).

Every new limb is a volume object in which each voxel contains a scalar representing the reconstructed signed distance. From this volume is possible to additionally extract a corresponding surface. Using this method we were able to create a time-course reconstruction of more complex object like a mouse limb with its corresponding flak (Figure 4A).

**Figure 4:**
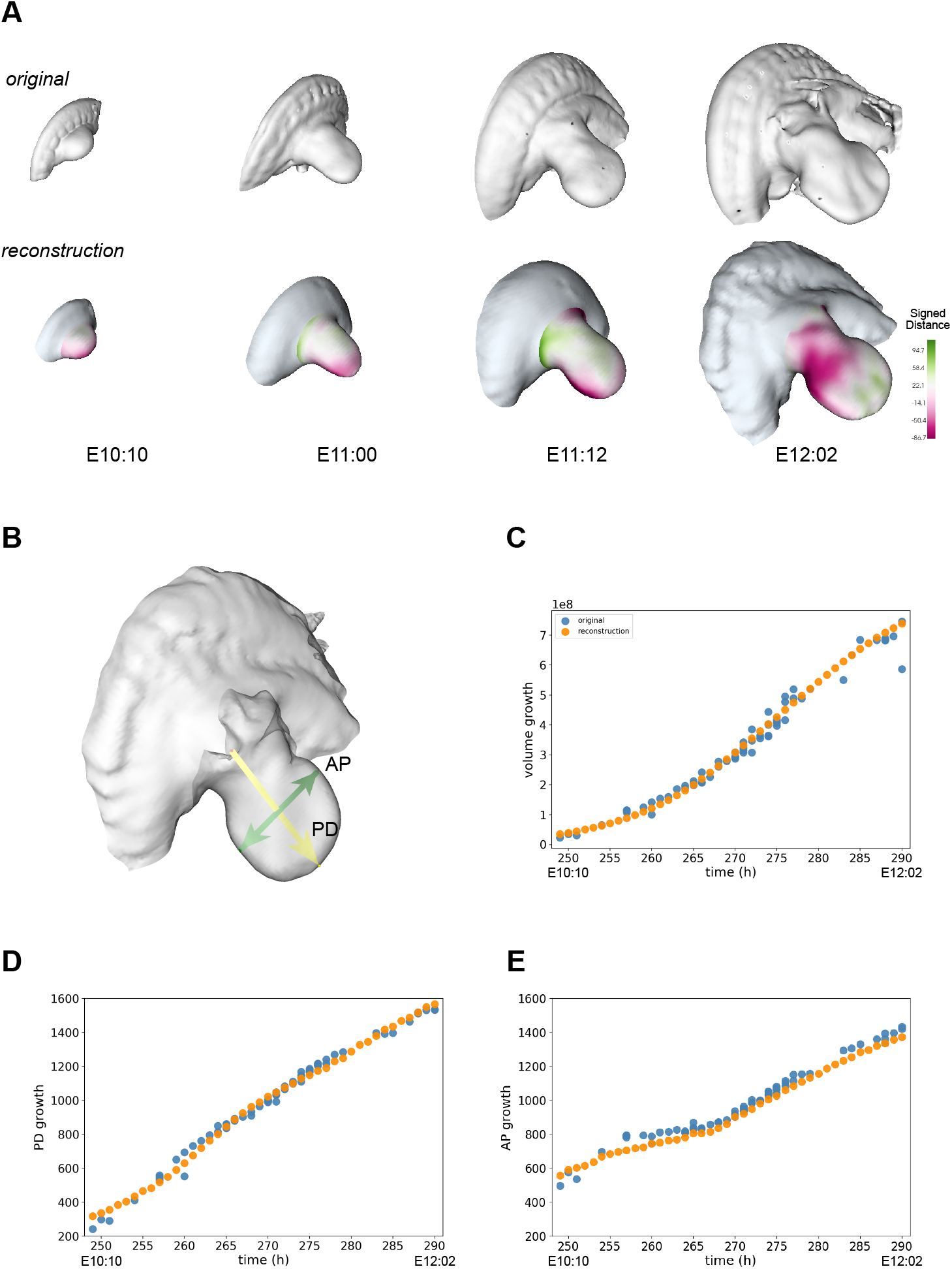
**A** - Comparison of original data (on top) at four different developmental stages (E10:10, E11:00, E11:12, E12:02) with the corresponding limb growth reconstructions (bottom), color map representing the signed distance between the reconstruction and the original surface. **B** - Representation of the proximodistal (PD) and anteroposterior (AP) axes of a limb. **C** - Volume growth comparison between original data (blue dots) and reconstructed shapes (orange dots). **D** - PD growth comparison between original data (blue dots) and reconstructed shapes (orange dots). **E** - AP growth comparison between original data (blue dots) and reconstructed shapes (orange dots).

The reconstruction in space and time of the growing limb consists of a smooth trajectory in developmental stages starting from E10:09 up to E12:02 (Figure 4A, Movie 2 - https://vedo.embl.es/fearless/#/limbflank). In order to know how accurately the result reproduces the 4D growth of a mouse limb, we directly compared it with the real data. To do so, we compared three distinctive characteristics of a mouse limb. Specifically, the total growth in volume of the limb, its elongation along the proximodistal (PD) axis and its widening along the anteroposterior (AP) axis (Figure 4B). For each reconstructed shape of a limb with its flank, the total volume was computed and compared with the volume of the original data at the same developmental stage (Figure 4C). In order to compare the limb elongation along the PD axis, we computed the geometrical distance between a fixed point (i.e. [0, 0, 0]) and the most distal point of the limb (Figure 4B) and we compared this value in the reconstructed and original data (Figure 4D). For confronting the enlargement of the limb along the AP axis the geometrical distance between the two furthest points in the AP plane of the limb “paddle” was determined and compared (Figure 4E).

The result represents the 4D-growth of an *ideal* limb which averages the common characteristics and features of all the limbs in the data set. In particular, we are recreating the growing process starting from when the limb bud is just a small bump of tissue (E10:09) and finishing when it already shows a distinctive “paddle” shape of the autopod (E12:02). Additionally, the reconstruction is able to capture the correct biological scaling of a growing limb by accurately reproducing the volume growth shown in the data and its proximodistal (PD) and anteroposterior (AP) axes evolution.

### 2.2 Application to the heart development

In order to determine the robustness of the method we applied it on a completely different set of data, specifically volumetric mouse heart data. We used a total of 26 OPT scans of mouse embryos where the heart is distinguished from the rest of the embryonic tissue by means of a molecular marker (myosin heavy chain MHC, as detected by antibody staining). The embryos were scanned by OPT and then processed to create surfaces representing only the MHC expressing part of the heart (Figure 5A). We obtained mouse heart data at six different developmental stages, i.e. 10, 14, 18-19, 21-22, 24-25 and 28-29 somites (Figure 5B).

**Figure 5:**
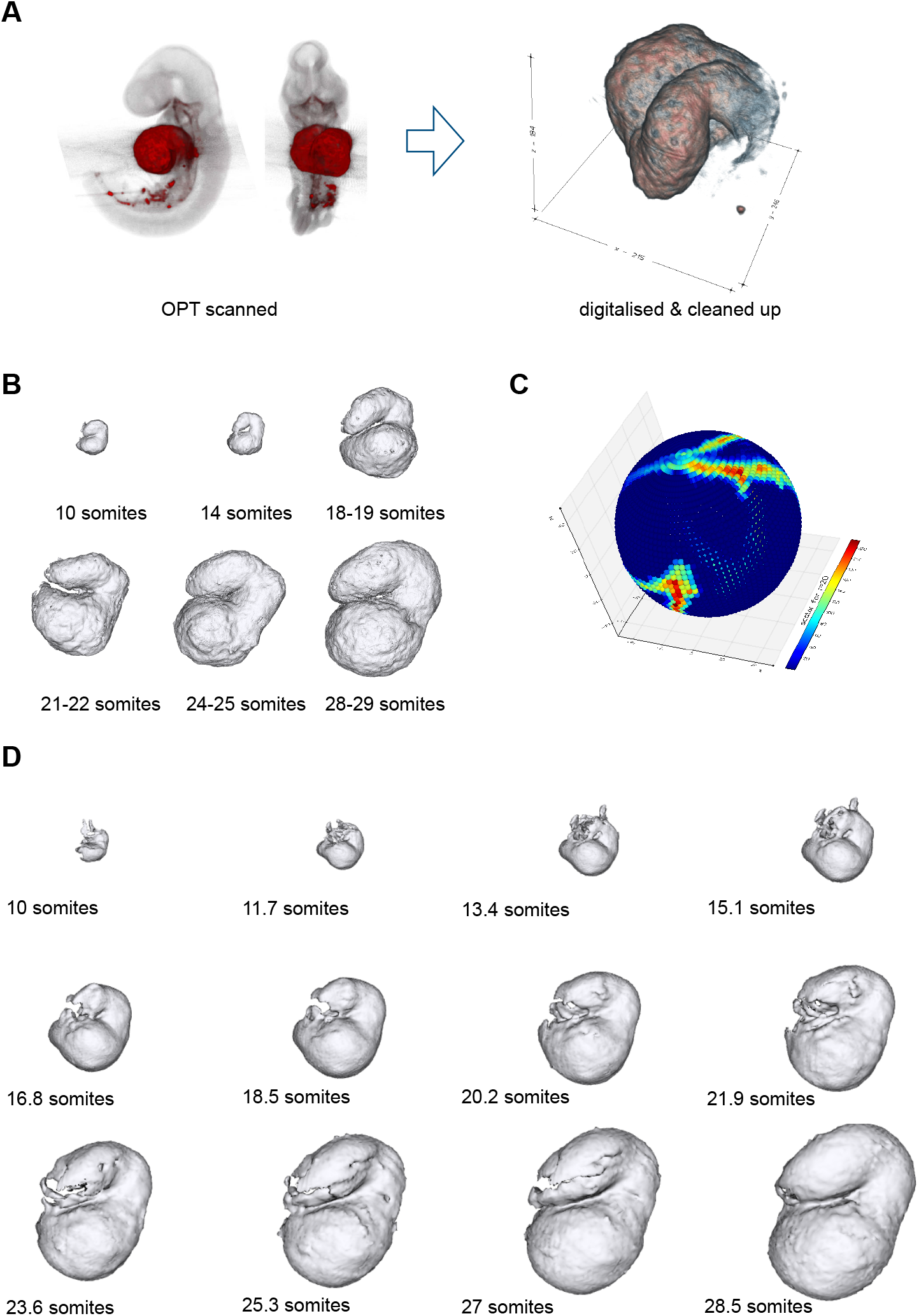
**A** - Representation of the data acquisition procedure: OPT scans of molecular marked mouse embryos, digitalisation and cleaning of the data. **B** - Examples of heart volumetric data at different developmental stages (i.e. 10, 14, 18-19, 21-22, 24-25 and 28-29 somites). **C** - Examples of scalar values of an 18-19 heart on one specific radius shell. Colors represent voxel intensities. **D** - The heart reconstruction at different developmental time points.

In this specific case, we considered the heart as a whole organ without the necessity of bending its surface. Therefore, differently from 2.1.1 it was possible to apply the method previously described directly on the volumetric data without the requirement of first converting them to surfaces and then using the signed distance.

As first step, all the data were manually aligned using a 4×4 linear transformation. Afterwards, similarly to 2.1.1, each heart was embed into a sphere centered at the geometric center of the object and the scalars, representing the voxel intensities, were computed along the radii of the sphere. The result is a set of intensities mapped on every concentric sphere from the radius discretisation (Figure 5C). The scalar values mapped on these spheres were expanded into spherical harmonics. The coefficients 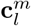 of the spherical harmonics were then interpolated over the developmental stages and these interpolated values were used to reconstruct the heart trajectory.

The results, as in the case of the mouse limbs (2.1.1), is a smooth trajectory of a growing heart in space and time that takes into account the common characteristics and properties of all the samples in the data set (Figure 5D, Movie 3 - https://vedo.embl.es/fearless/#/heart). Thus, in spite of the fact that mouse limbs and mouse hearts are markedly different in shape, our algorithm demonstrates to be completely general and to produce in both cases a reliable reconstruction of the evolution in space and time of the original data. Moreover, this approach, not only provides a quantitative basis for validating predictive models but it also increases our understanding of morphogenetic processes from a purely geometrical point of view.

## 3 Discussion

Our understanding of limb development (and other examples of organogenesis) has been driven by a strong focus on molecular and cellular activities. While modern biology has striven to make measurements of molecular data increasingly quantitative and complete (such as transcriptomics, and single-cell approaches), surprisingly, an accurate quantitative description of the continuously changing morphology of the limb has not been created. Precision in such a description has perhaps not previously been required for a basic appreciation of mutant phenotypes, but as biology becomes more predictive and quantitative, accurate measurements of dynamic morphology will be essential. For example, the value of quantification for static 3D limb bud shape has already been shown to increase sensitivity in phenotyping studies (Martínez-Abadías et al. 2018), as well as enabling the detection of subtle but real alterations in gene expression patterns. Quantifying dynamic shape changes will also become important for mechanistic and predictive models of morphogenesis. For example, we previously used data-driven finite element modelling (FEM) to rule out a popular hypothesis about limb bud elongation (Boehm et al. 2010). By combing quantification of the changing 3D shape with accurate measurements of proliferation, we were able show that the proliferation gradient hypothesis cannot be correct, because the model’s predictions do not fit the true morphology.

While for some model species a 4D quantitative trajectory can be captured by direct time-lapse imaging, this is not true for mammalian organs. In utero imaging cannot provide high spatial resolution, and in vitro culture techniques do not support normal organogenesis. We are therefore left with the challenge of reconstructing a 4D dynamic process, from static snapshots. Here we have developed and demonstrated a method to perform this task. It is based on the mathematical technique of spherical harmonics, which is a convenient method for dimensionality reduction of data that can be spatially distributed on a sphere. We were able to use the coefficients of the spherical harmonics from 69 different limb buds, spanning an age range of mE10:09 to mE12:02, and spline through these values to create the first smooth continuous 4D trajectory of normal mouse limb development. A simple, direct spherical harmonics method worked well on images just containing the limb bud itself, while a more complex volumetric version was required to interpolate a larger region of tissue that contained concavities (images which included a significant portion of the embryo trunk as well). A challenge for traditional landmark-based approaches, such as geometric morphometrics, is that the shape complexity dramatically increases from the beginning to the end of the trajectory, in other words presenting a changing amount of useful shape information. The spherical harmonics approach we present here copes well with this challenge, because of the intrinsic feature that coefficients of different orders (different levels of detail) contribute independently to the overall shape description.

In conclusion, we have applied this method to both limb development and heart development (the MHC expressing tissues) for the mouse embryo. It produces for the first time, a smooth, continuous and quantitative 4D description of their morphogenesis, including predictions for the gradual changes in lengths, and volumes of the tissue. We believe it can be applied to many other developing organs, and will be increasingly important as developmental biology becomes a more quantitative science, and moves towards predictive mechanistic computer modelling of morphogenesis.

## Methods

### Animals

All animal work was performed according to the guidelines of the Committees on Ethics and Animal Welfare established at PRBB and CNIC, in accordance to Spanish and European laws. C57Bl6/J mouse embryos were collected at the indicated gestational stages. For a precise staging of limb buds, the eMOSS staging system was used (Musy et al. 2018), and for hearts, pairs of somites were counted. Embryos were dissected in cold PBS and fixed overnight in 4% PFA at 4°C.

### Whole-mount antibody staining

Embryos were dehydrated with methanol and left overnight at −20°C. Whole-mount immunochemistry was performed according to standard protocols. Embryos were rehydrated, permeabilized in PBS plus 0.5% Triton X-100 (Sigma), left 2h in blocking solution (90% PBST (0.1% Triton X-100 in PBS)/10% normal goat serum) and incubated overnight at 4°C with anti-Myosin heavy chain, sarcomere (MHC) antibody (Developmental Studies Hybridoma Bank, MF20; 1:10). Embryos were washed in PBST four times for 4 hours and incubated overnight with biotin goat anti-mouse IgG (Jackson ImmunoResearch ref. 115-066-071; 1:500) and then with streptavidin-Cy3 (Jackson ImmunoResearch ref. 016-160-084; 1:500). After washing in PBST four times for four hours, embryos were stored in PBST 0.01% sodium azide and subjected to OPT scanning to obtain images.

### Optical projection tomography

Optical projection tomography (OPT) imaging (Sharpe et al. 2002) was used to acquire 2D images and obtain 3D reconstructions. MF20 immunostained samples were embedded in 1% low melting point agarose (Sigma), dehydrated in 100% methanol and cleared in BABB (1 volume benzyl alcohol : 2 volumes benzyl benzoate). Samples were scanned at intermediate resolution (512×512 pixels) in the Bioptonics OPT scanner using Skyscan software (Bioptonics, MRC Technology). The GFP filter (425/40nm, 475nm LP) was used to scan whole embryo anatomy, while the Cy3 filter (545/30nm, 610/75nm) was used to image the heart. OPT scans were reconstructed using NRecon software (SkyScan) and analyzed using the Bioptonics Viewer software.

### Implementation

The method is written in Python 3 and depends on standard python packages (Harris et al. 2020). All the mouse embryos were staged using the embryonic Mouse Ontogenetic Staging System (eMOSS) (Musy et al. 2018). Surfaces of the data were extracted from the OPT scans using the marching cubes algorithm (Lorensen & Cline 1987). The alignment in space of the mouse limbs was done using the reference provided by eMOSS and refined with the iterative closest point (ICP) algorithm (Besl & McKay 1992). Spherical harmonics transforms were made through the SHTools library (Wieczorek & Meschede 2018). The comparison between the reconstructed limbs and the original data was computed applying the signed distance using the angle weighted pseudonormal (Bærentzen & Aanæs 2005) implemented in *Vedo* (Musy et al. 2019). The bending of the flanks to make them match with the reference limbs was done by using a linear fit of *α* as a reference, and to apply a bending transform onto the shape data using *thin plate splines* (Bookstein 1989) (implemented in *Vedo* (Musy et al. 2019)), which is a non-linear transformation based on a physical analogy involving the bending of a rigid material. The limbs were not affected by this transformation, only the non aligned flanks were bent. All the visualisations and animations were produced using *Vedo* (Musy et al. 2019).

All the code is open-source and has been deposited in full in the GitHub repository → https://github.com/gioda/4D-reconstruction-of-developmental-trajectories-using-spherical-harmonics.

Additional movies, 3D interactive views of the results and further information can be found in the project’s web-page → https://vedo.embl.es/fearless.

## Acknowledgments

We thank Isaac Esteban and Miguel Torres for their invaluable discussions on shape comparison and spherical harmonics; Antoni Matyjaszkiewicz for his help with the web-page of the project. This research was supported by core funding from EMBL and CRG, and also the ERC Advanced grant SIMBIONT (project no. 670555) and a grant from the Fundacion La Marato TV3 (grant 082030). GD and MM were supported by the ERC Advanced grant SIMBIONT (project no. 670555).

## Author Contributions

GD: Conceptualization, Data curation, Formal analysis, Investigation, Methodology, Writing – original draft; MM: Investigation, Methodology, Writing – review and editing; MN, ARM, CBC: Performed the experiments; JSE Supervision, Funding acquisition, Writing – review and editing; JS Conceptualization, Supervision, Funding acquisition, Project administration, Writing – review and editing.

